# Damage explains function in spiking neural networks representing central pattern generator

**DOI:** 10.1101/2024.04.24.590949

**Authors:** Yuriy Pryyma, Sergiy Yakovenko

**Author notes:** Prepared for PLoS Computational Biology.

## Abstract

Complex biological systems evolved to control dynamics in the presence of noisy and often unpredictable inputs. The staple example is locomotor control, which is vital for survival. Control of locomotion results from interactions between multiple systems--from passive dynamics of inverted pendulum governing body motion to coupled neural oscillators that integrate predictive forward and sensory feedback signals. The neural dynamic computations are expressed in the rhythmogenic spinal network termed the central pattern generator (CPG). While a system of ordinary differential equations or a rate model is typically “good enough” to describe the CPG function, the computations performed by thousands of neurons in vertebrates are poorly understood. To study the distributed computations of a well-defined neural dynamic system, we developed a CPG model for gait expressed with the spiking neural networks (SNN). The SNN-CPG model faithfully recreated the input-output relationship of the rate model, describing the modulation of gait phase characteristics. The degradation of distributed computation within elements of the SNN-CPG model was further studied with “lesion” experiments. We found that lesioning flexor or extensor elements, with otherwise identical structural organization of reciprocal networks, affected differently the overall CPG computation. This result mimics experimental observations. Moreover, the increasing general excitability within the network can compensate for the loss of function after progressive lesions. This observation may explain the response to spinal stimulation and propose a novel theoretical framework for degraded computations and their applications within restorative technologies.

## Introduction

Skeletal dynamics of segmented body requires hierarchically organized neural control system with dynamic transformations expressed within spinal neural networks (Frigon et al., 2021; Yakovenko, 2011). The corresponding neural computations are thought to be embedded within the structure of nervous system. This is reflected within the topographical organization of motor cortex (Schieber, 2001). In the spinal cord, the motoneuronal organization embeds the musculoskeletal features of the controlled limb (Taitano et al., 2024; Yakovenko, 2004). The inverted pendulum mechanics describing limb interactions with the ground during locomotion (Kuo, 2007) is mirrored by the presence of spinal controllers with rhythmic pattern generation (Taga, 1995; Taga et al., 1991). The discovery of this neural ‘intrinsic factor’ or the central pattern generator (CPG) producing oscillatory output even in the absence of descending or sensory drives to control limbs in locomotion predates the formal description of the body dynamics (Saunders et al., 1953). Curiously, the first computational description of the CPG is one of the pioneering neuroscience models describing key dynamic mechanism within the nervous system (Verzár, 1923). Considerable number of both experimental and computational studies have examined this neural element over a century creating one of the most comprehensive theoretical descriptions of the nervous system that includes neuromechanical closed-loop dynamics with its control targets, the limbs. The study of disrupted computations in the CPG input-output transformation holds significant value for clinical applications for treatment of central and peripheral pathologies.

The application of the CPG concept in neurorehabilitation requires robust and efficient computational models that contain essential structure and function for a given behavior. The selection of the appropriate level of structural and functional details is imperative. On one hand, the Hodgkin-Huxley formalism has been used to capture ionic dynamics, and this method has proven to be useful for the understanding of oscillatory function of the CPG (Danner et al., 2017).

These models are typically designed to use large populations of neurons that are parameterized to recreate input-output relationships for the further theoretical post-hoc support of the experimental testing. The alternative and computationally transparent approach is to reduce the size of network to just several neurons, which reduces the model’s parameter space by several orders of magnitude. Surprisingly, these models can efficiently recreate the bursting characteristics underlying the locomotor pattern (Barnett and Cymbalyuk, 2014; Izhikevich, 2003). The simplification of the CPG with leaky-integrate-and-fire units can similarly capture temporal dynamics and even allow the real-time implementation for the control of neuromorphic simulations and robots (Angelidis et al., 2021). On the other hand, the simple models typically capture the high-level representation and can implement the broadly defined function of the CPG in regulating speed and the corresponding locomotor phases. Even relatively simple rate models can express the CPG output patterns without the spiking dynamics overcoming the parameter space problem and improving computational latencies in the input-output transformation (Sobinov and Yakovenko, 2018; Yakovenko et al., 2018, 2005).

The theoretical description of neural damage due to trauma or progressive degeneration is challenging. In this context, the drawback of simple neural models is their limitation in representing damage to their components—the neural substrate responsible for the neural computations. Spiking neural networks (SNN) offer a new type of formalism for CPG models that can capture the simplicity of rate models with the benefits of spiking neuron dynamics. In particular, machine learning tools developed for simple integrate-and-fire neurons offer a new biomimetic formulation that can be used in place of lumped-parameter rate models (Bekolay et al., 2014a). In this study, we have intended to accomplish two goals: 1) convert an analytical rate model of the CPG to the SNN implementation, and 2) to test the concept of impeded computations using damage within the network. Some preliminary results were published in abstract form (Pryyma and Yakovenko, 2022).

## Results

### Model validation

The CPG was formulated using two types of models, 1) a Brown-type rate model and 2) a network of simplified spiking neurons to validate the novel implementation, to study distributed dynamic neural computations and to test the contribution of functional identity to rhythmogenesis (Fig.1). The rate model for bilateral control of locomotor phases was previously developed in the context of extensor- and flexor-dominant patterns of rhythmogenic activity in cat fictive locomotion, a behavior observed in the reduced animal preparation with spinal rhythmogenesis and without all movement related feedback (Yakovenko et al., 2005). The function of the CPG rate model was recreated in the SNN-CPG model and validated with the experimentally defined temporal relationships governing the overground locomotion, as described below.

**Figure 1.**
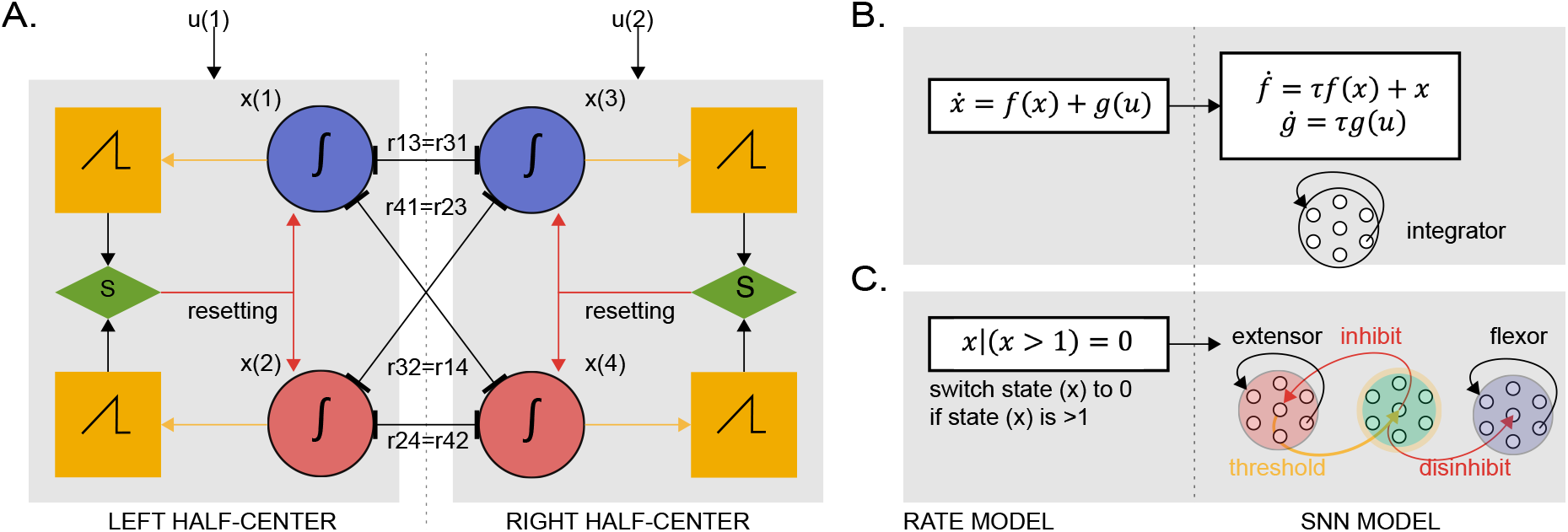
The oscillatory output is generated by two half-center oscillators with leaky integration and threshold resetting in the rate and SNN-CPG models. The half-center oscillator state (*S*) is set by the threshold crossing event (*x(j)≥0*) and ensures resetting between antagonistic signals, e.g., *x(1)* and *x(2)*. B. The integration process is represented by an ensemble of spiking neurons (*integrator*) expressing the changes in the internal state (*ḟ*) and external input (*ġ*). This dynamics of this system was embedded in the encoded neural state (Eliasmith, 2005). C. The switching from extensor to flexor state is triggered by threshold crossing of population output that initiates homologous inhibition and antagonist disinhibition.

The model performance was validated with the expected experimental relationship between locomotor phases and cycle duration (Halbertsma, 1983). A locomotor cycle is typically divided into stance and swing phases, corresponding to activation of flexor and extensor units of CPG. These experimental phase relationships can be simulated in the model with varied input into the CPG units (Fig. 2). The flexor phase is typically shorter than the extensor phase (Fig. 2A). The experimental relationships between the phases depend on the speed of walking. The extensor phase is more sensitive to changes in speed than the flexor phase (Fig. 2B, C). The examples in Figure 2 show representative performance of bilateral stepping pattern generated by three increasing levels of external inputs. The changes in flexor- and extensor-related states of left and right half-center oscillators are shown on the top and bottom plots in each column. The panels represent the simulation segments 6-10 s (A), 36-40 s (B), and 76-80 s (C) of a single simulation with a ramping up input (*u*). The reciprocal alternation of phases was generally symmetrical across both sides at across different speeds.

**Figure 2.**
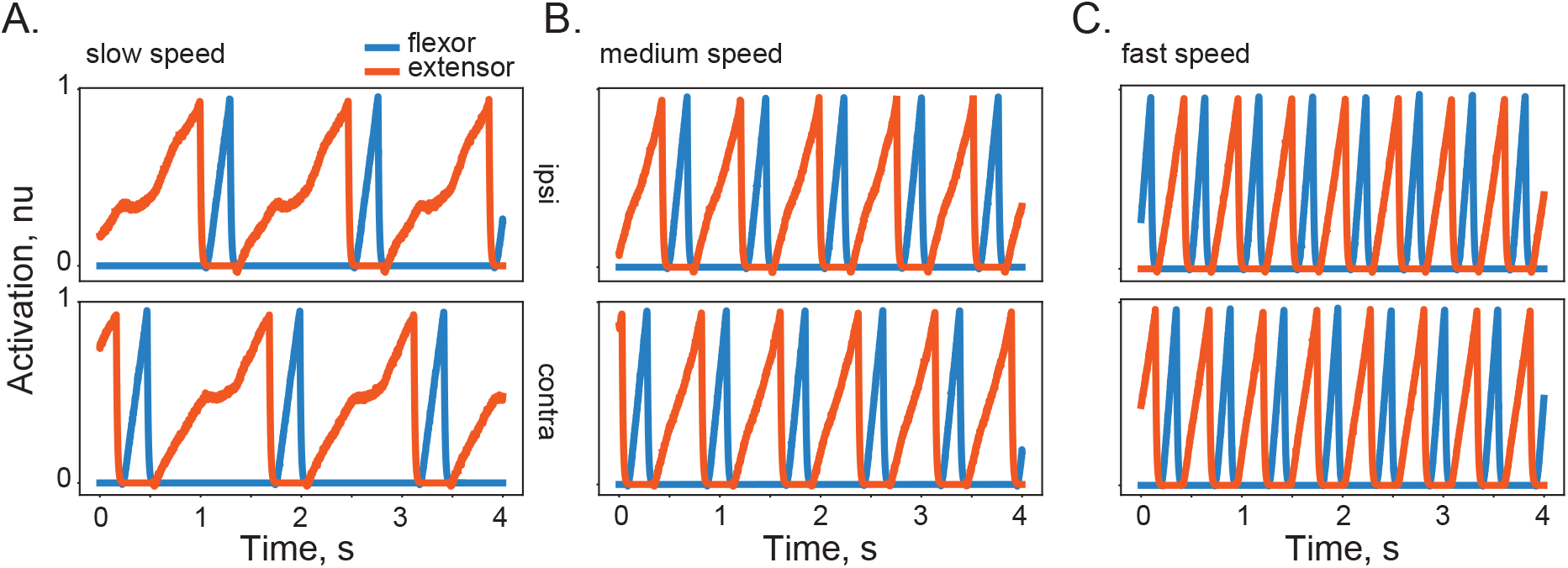
Examples of CPG model simulations with cycle periods representing slow, medium, and fast overground speeds. **(A-C).** The CPG inputs (*u* in Eq.1 in Methods) drive the dynamic transformation to generate the progression of system state defined with *ipsi* and *contra* pairs of half-center oscillators with reciprocal flexor and extensor signals. The activation values above zero represent the period of unit activity when it is integrating incoming signals and driving flexor or extensor motoneurons. The values equal to zero represent suppressed units and silent motoneurons.

The preferential modulation of stance (extensor) phase compared to swing (flexor) phase has been observed in multiple gait studies (Halbertsma, 1983), also seen in Figure 3A, black lines. The SNN-CPG model adequately captured the empirical phase relationships, reproducing the high gain and low offset of stance phase vs cycle period relationship and the opposite characteristics of swing phase vs cycle period relationship (Fig. 3A). The variation of cycle duration was achieved with a slowly changing input drive (*u* in Eq.1 in Methods). The simulated results recreated empirical measurements taken from previous study (Halbertsma, 1983) with high accuracy, R^2^=0.90 for swing and R^2^=0.99 for stance. the CPG input maps linearly on limb speed, a finding from the analytical solutions in the rate model (Sobinov and Yakovenko, 2018; Yakovenko et al., 2018). The same linear mapping between the input to the model and limb speed was observed for the SNN-CPG model (R^2^ = 0.99 with experimental results (Goslow et al., 1973); Fig. 3 B, C).

**Figure 3.**
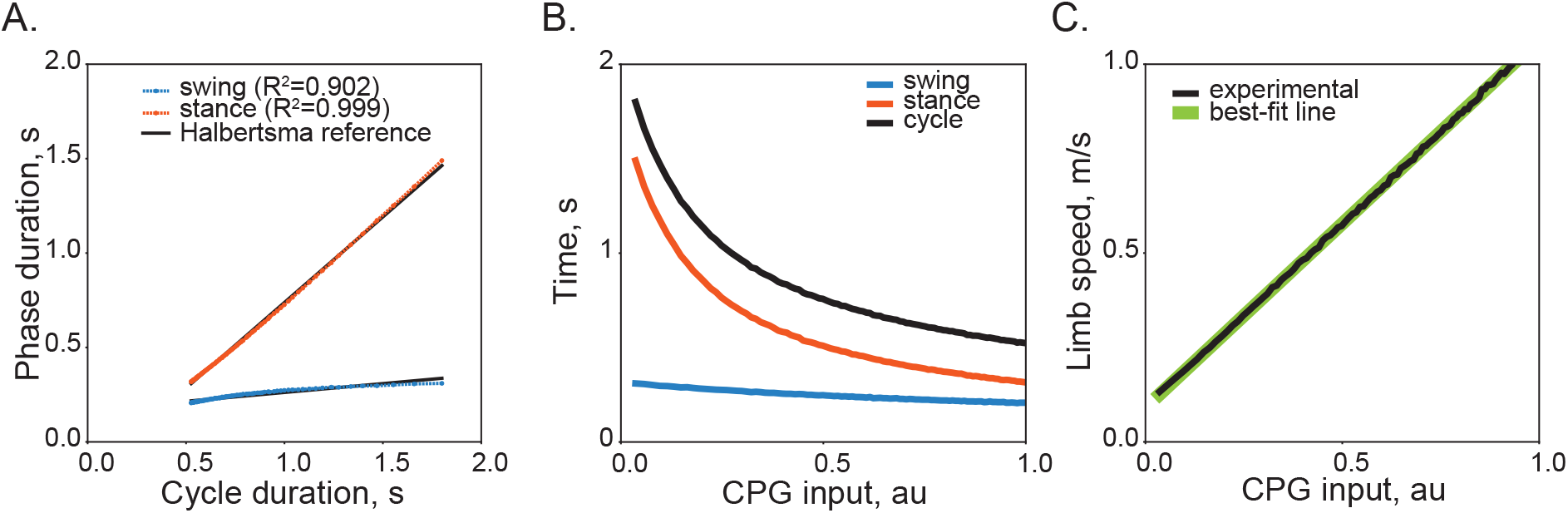
The SNN-CPG model validation through the temporal characteristics of phase modulation. **A**. The bilateral pattern of locomotor phase modulation with optimized SNN-CPG model parameters. The relationship between phase and cycle duration for each integrator is plotted together with the experimental relationship. **B**. The locomotor phase durations are related to the CPG input as a power function. **C**. The input in the SNN-CPG model is linearly related to the limb speed computed from experimental results (Goslow et al., 1973).

### Lesions and model performance

The substitution of rate equations with neural networks allowed us to test structural and functional correlates of neural damage and examine dynamic computations during progressive damage to its elements. The progressive damage to each of the four neural populations for left and right flexors and extensors led to different behavioral outcomes. We used the same cost function, previously used for the optimization of the model (see Methods, Eq. 6), to evaluate the computational error corresponding to the deterioration in the network with lesions. The error was expressed as the goodness of fit between phase and cycle durations, range, and bilateral symmetry. In a set of simulations with varying degrees of damage, the number of removed neurons ranged from 0.5 to 4.0 % of flexor, extensor, or both populations.

The error largely depended on the damage of extensor neurons (Fig. 4A), which is consistent with their function of encoding high gain and low offset in the phase vs cycle duration relationship (Fig. 3A). Examples in panels B and C illustrate this functional difference with two representative examples. The unilateral damage to the extensor unit disrupts the pattern but fails to terminate the ipsilateral extensor phase, skipping a step relative to the opposite (contralateral) side (Fig. 4B). This behavior is similar to the observations of so-called “flexor deletions” on the ipsi-side during fictive locomotion (Lafreniere-Roula and McCrea, 2005). In contrast, the pattern disruption caused by the equivalent flexor loss was not as detrimental as that caused by the extensor loss (Fig. 4C). The combined damage of flexors and extensors (Fig. 4A, black) generally followed the profile of extensor-only errors with only minor deviations at the highest level of damage.

**Figure 4.**
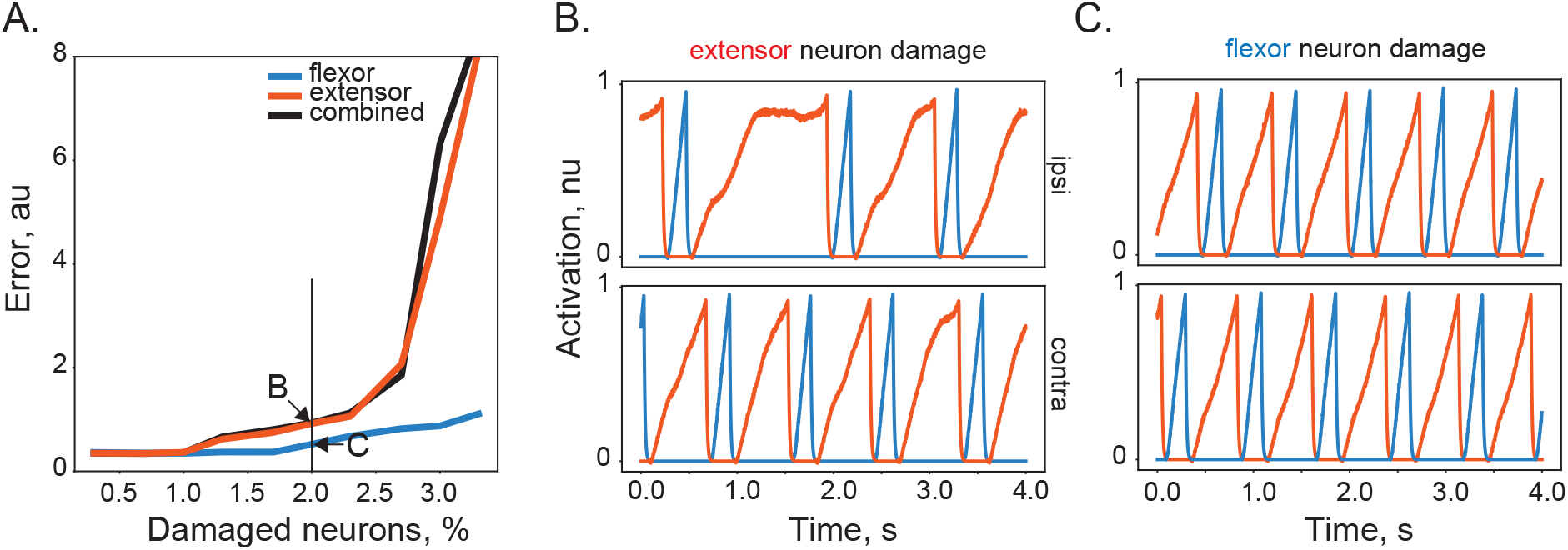
The damage of neurons in the SNN-CPG model leads to decreasing performance. The increase in model error (A) depends on the functional identity of “lesioned” neurons responsible for flexion (**blue**), extension (**red**), or both (**back**) phases. Examples of the output in models with 2% extensor (B) and flexor (C) ensemble damage.

### ‘Damaged’ computations and input drive

The concept of ‘damaged’ neural computations becomes tangible when the neural transformation is expressed through partial computations performed by an ensemble of tuned neurons. As our results have shown, the external drive to the system with damaged or missing components may no longer produce the desired outcome or accomplish intact function. The resilience of the dynamic CPG transformation was tested with the damage of varying volumes within the populations of either extensor or flexor neurons. To minimize the transient effects of initial conditions in the cost assessment, we started simulations without the disruption and applied the lesion after 5 s for 20 s. The relationship between the input drive, damage volume, and the SNN-CPG function was examined. Figure 5 shows the synthesis of results for extensor and flexor lesions. The heatmap combines sequential simulations for the various combinations of external inputs and neural damage that were iterated at 0.3% and 0.02%, respectively. The first phase error component (E_phase_) in Eq. 6 was used as the error value because these simulations were not exploring the full range of walking regimes. The area of low errors (yellow) denotes the network resilience to damage and is about four times smaller for extensors than for flexors, supporting the result shown in Figure 4. In both panels, the expansion of the low error area with the increase in the external drive indicates that the input drive to SNN-CPG rescues the locomotor pattern despite the neural damage.

**Figure 5.**
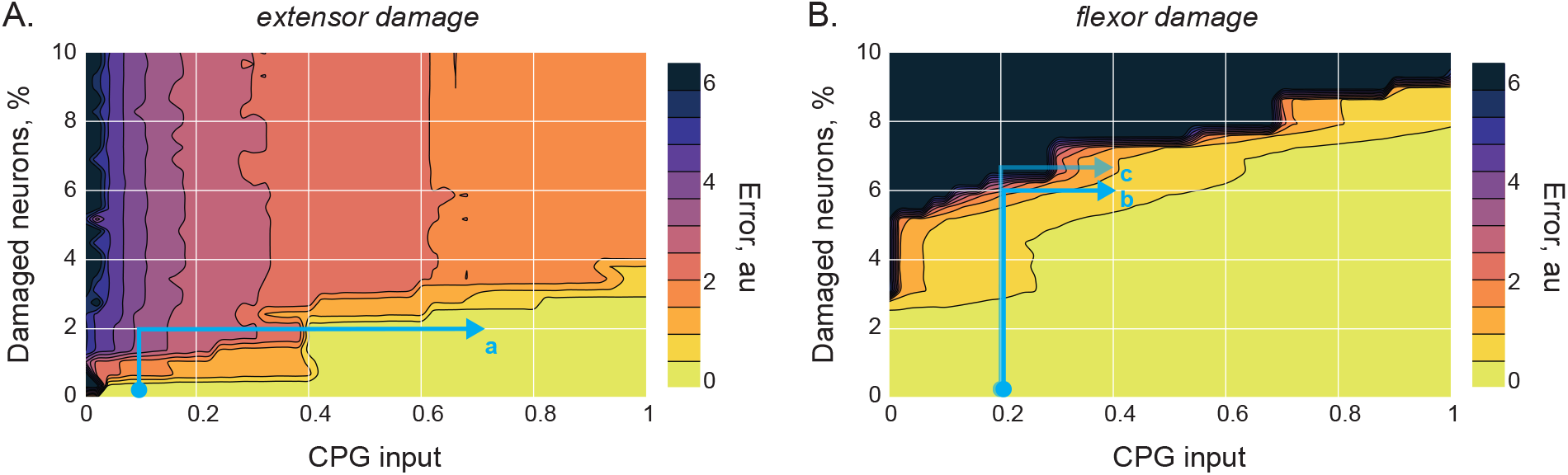
The relationship between the external drive, damage volume, and performance. The contour plots indicate changes in the quality of SNN-CPG in response to the damage to extensors (A) or flexors (B). The first component of cost function (*E_phase_* in Eq.6) was used as the error. Examples of simulations *a-c* are shown in Fig.6.

How does input drive rescue “damaged” computation? The examples of lesioned network patterns and their recovery are shown in Fig. 6. The simulation of extensor damage illustrates the custom pattern of simulation marked as (*a*) in Fig.5 A—the simulation starts with low input drive (0.1) and no damage but then is exposed to 2% damage at the same input (vertical blue line) and then rescued by the increased CPG input (0.7, horizontal line). Similarly, the simulation of flexor damage is shown with two examples in Fig.6 B&C following the patterns marked by trajectories (*b*) and (*c*) in Fig.5 B. These two simulations show the abrupt nature of the transition as the cost function is sensitive to one of the integrators locking into a given state as in the example of differences in 6% and 6.6% damage to flexors. The increase in the input drive rescues dynamic computations for both “damaged” populations.

**Figure 6.**
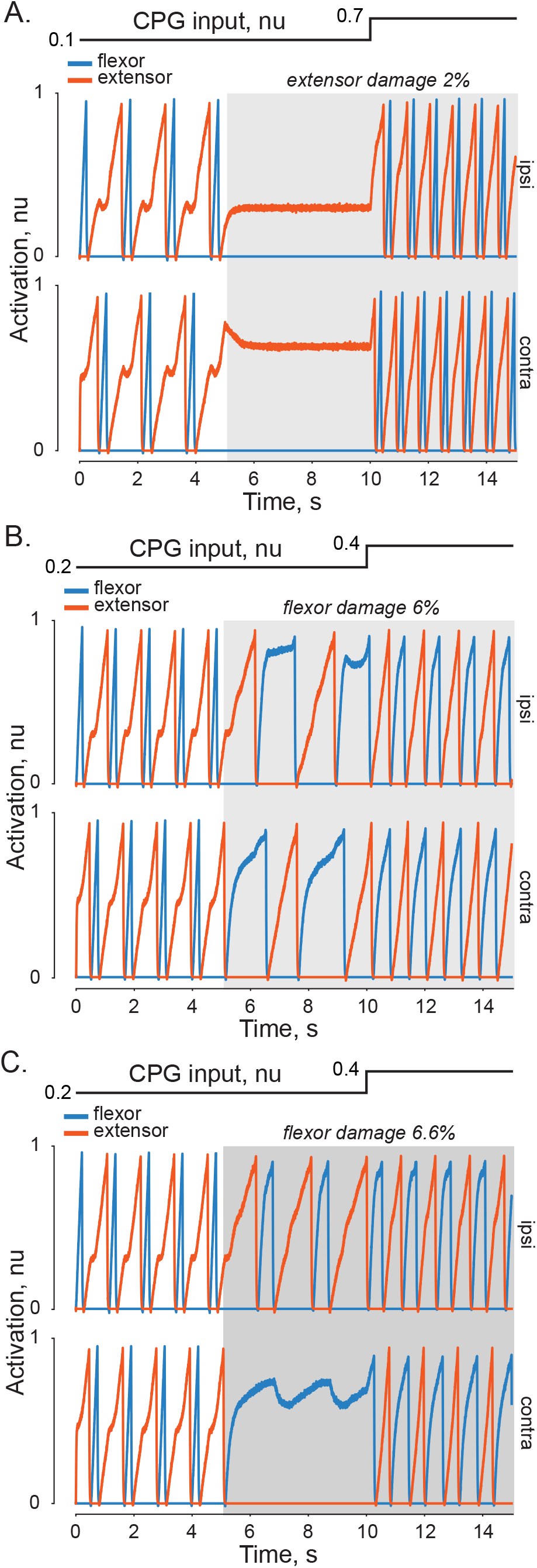
The examples of extensor and flexor damage with different drive levels. A. The bilateral pattern of ipsi- and contra-half-center dynamics is shown for the custom pattern in Fig.5A (*a*). The simulation with the low drive without damage followed by 2% damage to extensors at the same (0.1) and then at the increased level of external drive (0.7). B&C panels similarly show two examples of flexor damage described by the custom simulation patterns (*b*) and (*c*) in Fig.5 B.

## Discussion

In this study, we expressed CPG structure and function within a network of artificial spiking neurons and examined the effect of simulated progressive damage on functional dynamics. We used networks of leaky-integrate-and-fire neurons trained to embed the rhythmogenic function of the spinal locomotor CPG previously described with the rate model. The neural network replicated the accuracy of our relatively simple rate model, which is evident from the same input-output relationships in Fig. 3. Similar to our previous studies (Sobinov and Yakovenko, 2018; Yakovenko, 2011; Yakovenko et al., 2018), this model reproduced the regulation of swing and stance phases and maintained the linear relationship between the input drive and stepping speed. Similarly, the correlated relationship between the oscillation frequency and input drive have been confirmed in the models with Hodgkin-Huxley formalism (Danner et al., 2016). While the dynamical profile of the input drive was not predefined, the increase in the drive changed gait from walk to trot, gallop, and bound. The linear relationship between the spinal activation and output speed was computed from cycle durations (Goslow et al., 1973) in the models of CPG describing extensor- and flexor-dominated patterns in fictive locomotion (Yakovenko et al., 2005) and supported by neuromechanical overground gait simulations (Yakovenko et al., 2004). Moreover, the afferent locomotor drive was also shown to interact with the CPG drive and compensate, if needed, for the discrepancy between the CPG oscillation and the mechanical state. This neuromechanical tuning theory provides the common basis for the interactions between predictive and adaptive processes (Prochazka and Yakovenko, 2007).

Historically, lesions at different levels of neural hierarchy have traditionally been one of the main neuroscience tools for studying function and structure with complex interactions. The prominent examples are the systematic lesioning studies of the corticospinal and brainstem pathways in monkeys (Lemon et al., 2012). Another example is the discovery of dorsal and ventral stream processing or “where/how” and “what” visual processing pathways (reviewed in Milner and Goodale, 2008). Numerous studies of action and perception representations using recording, stimulation, and lesion methodologies refined the view of differences in these cortical computations for distinct functions of visual processing. Further down the neuraxis, spinal lesions in vertebrates provided the foundation for the understanding of spinal processing (Sherrington, 1906) and intrinsic spinal rhythmogenesis (Brown, 1911). The classic work was followed by the identification of structural elements involved in rhythmogenesis showing colocalization of rhythmogenic circuitry with the motoneuronal targets (Kiehn and Kjaerulff, 1998; Rossignol et al., 2002, 1996). Unsurprisingly, the spinal lesions caused the decrease in the excitability and, subsequently, the computations of spinal locomotor and postural pathways (Mari et al., 2024; Zelenin et al., 2016) that could be restored with the electrical stimulation below the injury (Yakovenko et al., 2007). In this study, we simulated controlled and specific lesions to understand the relationship between structure and function of separate populations of simulated neurons within the CPG network. Our result is the demonstration of functional differences in flexor and extensor networks using progressive injury in the model. Similar experimental methodology was used to understand “deletions” in fictive locomotor output resulting in the CPG models with separate networks for rhythm generation and pattern formation computations (Lafreniere-Roula and McCrea, 2005; Mccrea and Rybak, 2008; Perez et al., 2009; Rybak et al., 2006).

Using lesions within the SNN model, we discovered that the trained functional identity of neurons was differentially affecting the overall CPG computation. Figures 4 and 5 show that the loss of a small neural volume of extensor networks reduced the coordination and phase regulation much more than the equivalent flexor volume loss; however, the degradation of flexors had a rapid transition to the disrupted pattern. The relatively higher velocity-dependent modulation of extensors than that of flexors indicates that damage to the extensor-related pathways, playing a gravity-related role during overground locomotion, has a higher impact on the state of the CPG. However, in fictive locomotion induced by stimulation of the mesencephalic locomotor region in animals with paralyzed muscles and no movement-related sensory feedback or extensor gravity-related drive to the CPG, the bias can switch to the flexor-dominated pattern instead (Yakovenko et al., 2005). We speculate that the damage to spinal flexor pathways during this state with flexor bias would have a higher impact on the overall CPG function than the damage to extensor pathways, which is the opposite of what we observe during the overground locomotion. Overall, these findings support the idea that the contribution of structurally identical neural networks may differently impact neural computation according to the embedded function and dynamic regime of the system.

The concept of “damaged” neural computation is novel in the context of movement control. Previously, a similar approach of simulated injury in a neural network was applied to the study of aphasic naming from the recovery dynamics (Wright and Ahmad, 1997). The dropout regularization technique is a standard approach that randomly removes subsets of neurons during training to control overfitting (Wager et al., 2013) and allowed us to resolve training of the posture-dependent musculoskeletal transformations (Smirnov et al., 2021). Generally, the outcome of structural manipulations is difficult to assess in models with single neuron and network dynamics. Even a simple spiking neuron model can generate a dozen of possible regimes, including the CPG-like bursting (Izhikevich, 2007). The expression of the same function by neural networks with varied structure and composition is seen in diverse neural ensembles of bursting neurons reproducing the alternating neural pattern (Prinz et al., 2004).

In our modeling approach, we holistically express the CPG function within the structure and dynamics of a neural circuit. Starting with one of the simplest mathematical expressions of the locomotor CPG, we converted its half-center dynamics from rate functions to be expressed within an ensemble of spiking neurons implemented with a range of parameters to accomplish the distributed computation. Each population integrated inputs in the presence of interactions with other populations. When a portion of neurons was virtually lesioned in each population, the overall dynamical transformation of desired limb speed to appropriate locomotor phases was gradually disrupted. The quality of output could be restored with the additional input as demonstrated by the parameter space exploration in Fig. 5. Because this model is relatively transparent, this finding can be expected. An analogy of this *damaged computation* is depicted as the ‘*rusty watermill*’ mechanism in Fig. 7. The input-output transformation is a rotation powered by the force of water and gravity (an overshot wheel) to drive axial rotation. The rotation can be impeded by the “rusty” axel, but additional input drive can overcome the lost torque due to friction to achieve the expected rotation dynamics.

**Figure 7.**
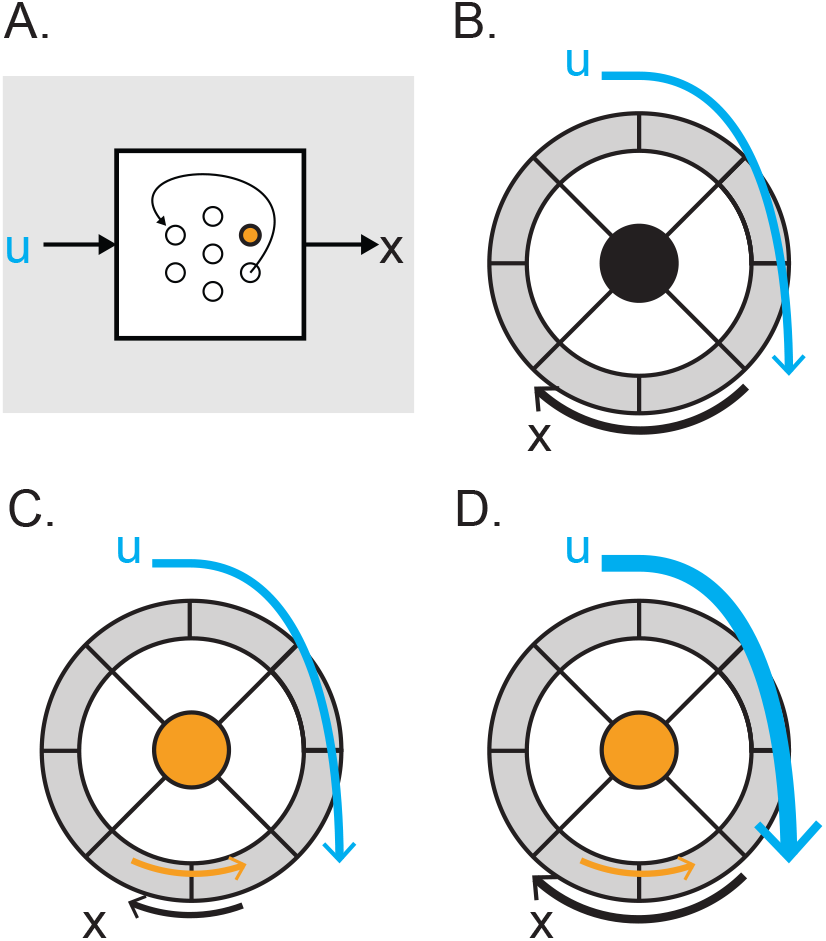
The concept of impeded ‘rusty watermill’ computations. A. The input-output transformation depends on computations performed by units within SNN. B. The hypothetical “watermill” computation transforms input (u) into the rotation moment (x). C. The reduced computational function of “damaged” neurons (orange) can be analogous to the decreased generation of the rotational moment in the concept model. D. The increased input can restore the desired movement/computation.

This simple thought experiment of the *damaged computation* provides a simulation of the impeded integration process that can provide intuitive predictions for the experimental observations. For example, epidural electrical stimulation of damaged spinal cord can restore rhythmic limb movement even with a continuous and nonspecific exogenous input to the spinal pathways in humans (Harkema et al., 2011). The nonspecific electrical stimulation of descending propriospinal pathways above the lumbar enlargement can also improve rhythmogenesis even in animals with a complete spinal cord transaction (Yakovenko et al., 2007). In general, increasing basal excitation of efferent and/or afferent circuits leads to the recovery after spinal cord injury (reviewed in Eisdorfer et al., 2020). These outcomes of epidural electrical stimulation are consistent with our model predictions where the network computation can recover with the increased external drive from either afferent or efferent circuits, as illustrated by examples in Fig. 6. In these simulations, the “injury” typically slowed the frequency of oscillations (locomotor speed) predicting that injury severity correlates with the rate of neural dynamics. This is generally supported by observations of correlation between the reduction of locomotor speed and progressive neurodegenerative disorders. The locomotor speed helps guide clinical management and rehabilitation strategies (Graham et al., 2008; Middleton et al., 2015).

There are several potential limitations in this model. First, the neural dynamics of CPG does not include the coordination with the mechanical system. In neuromechanical models with both pattern generation and afferent feedback, the lack of central drive can be compensated for by the increase in the gain of sensory feedback (Di Russo et al., 2023; Markin et al., 2010; Yakovenko et al., 2004). Also, adding or removing stretch reflexes during locomotion can modulate the limb speed (Prochazka et al., 2002; Yakovenko et al., 2004), which supports the idea that a common modality such as limb speed is used for the integration of multiple feedforward and feedback signals.

Second, the CPG model is based on the classical Brown’s model of half-center oscillators (Brown, 1911; Verzár, 1923) without the separation into the rhythm generation and pattern formation layers (Rybak et al., 2006). The support for the disparate populations of spinal rhythm and pattern-generating interneurons comes from the observations of “deletions”, the interruptions in the pattern of one of the functional groups with reinstatement of the rhythm in the following cycles (Lafreniere-Roula and McCrea, 2005), which can also be observed in our examples Fig. 4B and Fig. 6C. In our model, the cycle timing is preserved as a network dynamic property. This demonstration shows that the observation of “deletions” does not imply that the network requires the separation of rhythm and pattern-generating layers. Indeed, given that the simplest models of spiking neurons have regimes of intrinsic bursting (Izhikevich, 2003), many types, if not all, biological neurons may operate within a rhythmogenic state. Even the spinal motoneurons can be endogenously rhythmogenic as observed in a functional continuum of rhythmogenesis among zebrafish motoneurons (Menelaou and McLean, 2012) or participation of motoneurons in modulating phase durations seen in leech crawling (Rotstein et al., 2017). When describing the rhythmogenesis of CPG, it is challenging to dissociate the contributions of endogenous and network mechanism of rhythmogenesis among both spinal interneurons and the co-localized motoneurons (Barkan and Zornik, 2019). The bistable firing behavior of vertebrate motoneurons has been observed readily in injured and intact vertebrate neural system showing hysteretic and facilitation dynamics in response to repeated inputs (Binder et al., 2020; Kiehn and Eken, 1997). Overall, these observations support the idea that many spinal neurons and even output layer neurons (motoneurons) may contribute to rhythm generation; the disruption of activity within a subpopulation of this network may not indicate that the missing part was responsible only for the pattern formation function. Thus, the separation of the population into rhythm generating and pattern formatting networks may not be essential and could theoretically arise from the continuum in the distribution of parameters that are responsible for endogenous activity.

Third, the size of each network was heuristically defined. About 300 neurons generated smooth representations of state dynamics with diminishing gains in further upscaling of the population size. This number of neurons is similar to that in another study using SNNs to describe the CPG controlling neuromechanics of a lamprey, where about 280 – 2000 neurons were used to recreate the segmental coordination (Angelidis et al., 2021). It is speculative to draw parallels to the similar size of motoneuron populations in vertebrate spinal cord since simulated neurons include simplistic spike generation dynamics, oversimplified structural complexity, and do not reflect unit type representation or the detailed organization observed in the spinal cord (Taitano et al., 2024; Yakovenko et al., 2002). The digitization of computations performed by our original rate equations and the temporal resolution of desired output are likely reflected in the size of population encoding the input-output transformation.

In conclusion, we report a novel finding that the functional identity of flexor- and extensor-related neurons in the CPG differentially affects the dynamic computation of this mechanism. This was revealed by the different system responses to the progressive “lesions” of flexor and extensor SNN populations. In addition, the CPG input increasing the general excitability within the network can compensate for the loss of function after damage, restoring the original neural computation. Moreover, the simulated result of damage within the dynamic computation of the CPG network indicates that the separation into rhythm generation and pattern formation layers is not essential for the description of “deletions” observed within the fictive locomotion pattern. The modeling of the CPG mechanisms with SNNs offers insights into the neural dynamics of spinal rhythmogenesis and identifies therapeutic targets for restoration.

## Methods

The development of the locomotor CPG model using SNN formalism was based on the conversion of the previously developed rate model implemented as a system of ODEs with leaky integrations. Rather than using the full implementation of Hodgkin-Huxley formalism, we chose to expand the implementation to the simplified models of spike generation. The novel implementation required the conversion of the following components: i) the representation of an artificial neuron network with distinct generally reciprocal states; ii) the transition dynamics between bursting and non-bursting states; iii) the implementation of objectives for model training and validation.

### Model structure

The model of bilateral flexor-extensor circuit of CPG (Fig. 1A) was simulated by a system of four leaky integrator equations constituting the rate model of CPG (Yakovenko, 2011). In this study, this model was re-expressed using the network of the leaky integrate-and-fire (LIF) neurons with simple spiking and tuning dynamics (Bekolay et al., 2014b). The rate model state consisted of four scalar variables *x* = [*x*_1_, *x*_2_, *x*_3_, *x*_4_]^*T*^ describing the state of bilateral flexor-extensor networks. The dynamics of this model has been previously described in the context of capturing phase dominance phenomenon observed in fictive locomotion (Yakovenko et al., 2005), limb control (Sobinov and Yakovenko, 2018), and interlimb coordination during turning (Yakovenko et al., 2018). Similar to our previous work, the model is described by the leaky integration of external inputs (*u*) and internal states (*x*) in Eq. 1, where the contributions are parameterized with the corresponding gain matrices (*Gu*, *Gx* unilateral and bilateral gains) with individual parameters (*r*_*ij*_) described in Eq.2-3:

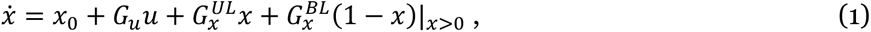

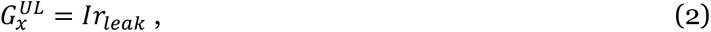

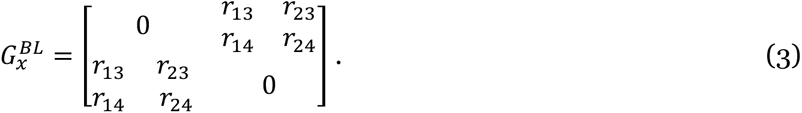

Here, the indices *i* and *j* correspond to the connectivity structure in Figure 1 between the half-centers, where flexors are (1) and (3) and extensors are (2) and (4). Symmetric parameters were set the same values as shown in Figure 1A to reduce the parametric space.

Similarly, we represented four states with four neuron ensembles of neurons simulated with a simple LIF neuron model. This model is built on the simplified principles of spike generation where ionic channel dynamics is governed largely by [Na^+^] current triggering of action potential and with the automatic resetting of membrane potential after spike generation. Thus, the spiking frequency of a neural population can be encoded to embed an arbitrary input-output relationship. We used Nengo, which is the framework for the development of custom structural networks expressing an embedded input-output function (Stewart et al., 2009). In brief, the process of encoding is an error-driven adjustment of spike generation related parameters for each neuron describing its tuning curve, which is inspired by the population coding (Georgopoulos et al., 1986). All parameters related to spike generation are the neuron gain (*⍺*_*i*_), its baseline firing rate 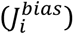, and the encoding weight (*e*_*i*_), but only the encoding weight parameter was used in the error-driven encoding of population tuning in this model. Neuron encoding parameters were sampled from uniform distributions (maximum rates from [200, 400] and intercepts from [−1.0, 0.9]). Parameters *⍺*_*i*_ and *e*_*i*_ were solved from maximum rates and intercepts to satisfy LIF neuron properties. The encoded state was applied through recurrent connections to all neurons within each simulated ensemble of N neurons (Fig. 1B). The spike train output of each neuron was the transformation of spike generation parameters through a LIF function (*G_i_*), as described in Eq.4:

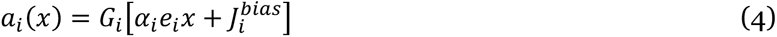

The decoding process of computing the estimated current state (**x̂*_J_*) was the summation of all filtered spike trains multiplied by their decoder weights (*d_i_*). These weights were determined by solving the least-squares problem with **x̂*_J_* = *x_j_*:

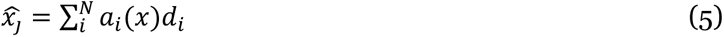

In our model four (*j* ∈ [1,4]) neural populations (N_j_=300) encoded four states (*x*) representing bilateral locomotor phases. The corresponding representation error for the state variable–the difference between encoded and desired output–was within noise tolerance (average RMS error of 0.003). The output of each neural ensemble ensured appropriate switching between locomotor phases. Similar to the rate model, the thresholding of outputs determined phase switching using the LIF threshold neuron (orange box in Fig.1A) defined on the range [−1, 1]. During encoding, the stochastic integrator output (**x̂*_J_*) and the threshold neuron spiked when input exceeded the intercept (set to 0.9 with sampling rate of 400). Next, the “switch” ensemble of LIF neurons (green box, N=2, intercept 0.0, sampling rate 400) encoded the *resetting* output of either −1 or 1 to enforce the reciprocal relationship between flexor-extensor half-centers (1-2 and 3-4 in Fig. 1A). The output of −1 inhibited extensor ensembles by setting the encoding weights of the extensor population to 0 so that any input would be unable to generate spikes. Similarly, the output of 1 inhibited the flexor ensemble.

### Simulations and cost function

The representation of flexor-extensor phase characteristics was achieved through the minimization of an analytical cost function in continuous simulations with a span of continuously increasing inputs (u). The cost was expressed as the scalar error relative to the target intra- and inter-limb phase characteristics. Similar to our previous work, the objective function consisted of the following components: 1) intralimb phase duration error; 2) interlimb coordination error; 3) intralimb cycle range error; and 4) interlimb symmetry error:

1. The intralimb phase duration error (*E*_*phase*_) was calculated as the root-mean squared (RMS) of the target and simulated phase values. The target values were expressed as the analytical best-fit functions (Halbertsma, 1983). The simulated flexor and extensor values were detected in the corresponding simulated outputs through thresholding (*S>0* switches to stance and *S<0* switches to swing). The cycle duration for each detected step was calculated as the sum of two phases.
2. The interlimb coordination error (*E*_*coordination*_) discouraged the simultaneous flexion phases in both limbs. Typically, a swing phase in one limb is supported by a stance (support) phase in the contralateral limb during overground walking.
3. The intralimb cycle range error (*E*_*ran*g*e*_) encouraged the parametric solutions that generated the expected experimental range of cycle durations.
4. The interlimb symmetry error (*E*_*symmetry*_) promoted symmetrical occurrence of flexion phase in the middle of the contralateral extension phase.

The corresponding errors were calculated separately for each limb and summed. The total error (*E*) was a sum of all components scaled by custom coefficients reflecting subjective importance of each error source:

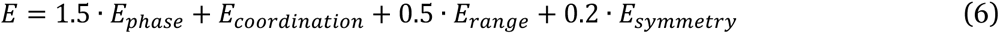

### Network optimization

SNN model optimization has two algorithmic steps: 1) the optimization of CPG model parameters and 2) the solving of decoding weights that minimize the representation error with these model parameters. We used the system of ODEs from Eq. 1 to describe the input connections to each integrator *x_j_* with optimized parameters *r_ij_*, *r_leak_*, *G_u_*, *x_0j_*, defined on the ranges described in Table 1 and then adjusted sequentially based on the error in the output (Eq.6) using hyperout method (Kim et al., 2018). The hyperout method was faster than the Nelder–Mead method (MATLAB, The MathWorks Inc.). The following example of this process based on Eq. 1 describes the rate of one of the half-centers in the model (*ẋ*_1_):

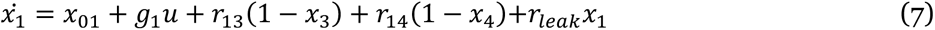

**Table 1.** Explored range of model parameters. All parameters are from Eqs. 1-3. The initial condition of flexors (*x_ic1_*) was set to 0.0 and extensors (*x_ic2_*) was set as a search parameter. The optimization minimizing the cost function determined a set of optimal values from the min-max range.

| Variable | $x_{o1}$ | $x_{o2}$ | $g_1$ | $g_2$ | $r_{leak}$ | $r_{13}$ | $r_{14}$ | $r_{23}$ | $r_{24}$ | $x_{ic2}$ |
| --- | --- | --- | --- | --- | --- | --- | --- | --- | --- | --- |
| **min** | 4 | 0 | 2 | 2 | -1 | -1 | -1 | -1 | -1 | 0 |
| **max** | 6 | 2 | 5 | 5 | 1 | 1 | 1 | 1 | 1 | 1 |
| **optimized** | 4.6773 | 0.5019 | 3.4240 | 3.8701 | 0.4442 | 0.0827 | -0.9559 | -0.9528 | 0.7736 | 0.4960 |

These optimized parameters for all integrators (*r_13_*, *r_14_*, *r_leak_*, *g_1_*, *x_01_* for integrator *x_1_*) were locked during the test simulation. The error in the output was evaluated using the cost function for 80 s of model simulation with the input signals (*u*) linearly growing from 0 to 1. The cost was computed from the simulated swing and stance phases for two limbs using Eq.6. Table 1 shows the optimized values corresponding to the minimal cost. The spiking train was generated by the network expressing the vector (*x̂*) described Eq. 5.

### Simulated damage

The formulation of CPG dynamics with spiking neurons enabled us to examine the effect of progressive damage in simulated “lesion” experiments. The first hypothesis tested here was that function can define relative importance of different cell groups participating in locomotor pattern generation. Specifically, we removed structurally identical simulated neurons by assigning zero to a select proportion of either *flexor, extensor,* or *both* neurons and compared the resulting change in cost. The change in the performance error was evaluated in a set of 80s simulations with the removal of 1 to 10% of cells in specific neural populations. The second tested hypothesis was that the external drive can rescue the damaged neural computation. The different types of neural lesions were imposed on the model followed by the ramp of increasing external drive. The ramp from 0 to 1 was applied for 20s. In both tests, the cost was calculated according to Eq. 6.

## Acknowledgements

We would like to thank Valeriya Gritsenko for the helpful comments on this manuscript.

